# Toads and roads: transport infrastructure severely impacts dispersal regardless of life stage

**DOI:** 10.1101/349639

**Authors:** Hugo Cayuela, Éric Bonnaire, Aurélien Besnard

## Abstract

Transport infrastructure such as roads has been reported to negatively affect dispersal. Their effects on dispersal are thought to be complex, depending on the characteristics of the structure and the intensity of the traffic using it. In addition, individual factors, such as age, may strongly affect dispersal decisions and success when individuals are confronted with transport infrastructure. Despite the importance of this topic for wildlife conservation, few studies have investigated the effect of transport infrastructure on individuals’ dispersal decisions before and after sexual maturity. We examined the effects on two kinds of infrastructure, gravel tracks and paved roads, on the dispersal of an endangered amphibian, the yellow-bellied toad (*Bombina variegata*). We used capture-recapture data collected during a five-year period on a large, spatially structured population of *B. variegata.* Our study revealed that emigration rates increased with an individual’s age, while dispersal distance decreased. It also showed that both tracks and roads had negative effects on dispersal. The negative effect of roads was stronger than that of tracks. We additionally found that the effect of tracks on dispersal slightly decreased with a toad’s age. In contrast, the negative effect of roads was severe and relatively similar across age classes.

## Introduction

The impact of transport infrastructure (i.e. gravel tracks, primary and secondary roads and railway lines; hereafter, TI) on biodiversity has received considerable attention from ecologists as well as conservationists over the last three decades^1–4^. In industrialized countries, the intensive development of transport infrastructure since the 1970s has had severe impacts on biodiversity^2,5^. Four main ecological impacts with synergic effects are usually highlighted (see references above). First, the construction of TI always triggers a net loss of natural habitats, resulting in population declines and local extinctions. Second, collision with vehicles can cause mortality in organisms crossing TI, which may lead to massive mortality and population decline. Third, both the use (i.e. traffic) and maintenance of the infrastructure (e.g. sanding or salting icy roads) alter and pollute the habitats located in the direct vicinity, which can durably affect the biology of organisms (e.g. physiology and behavior), population dynamics and species distribution. Fourth, TI may either limit or facilitate movement, thus acting as a barrier or a corridor, which can profoundly affect the dynamics of spatially structured populations.

Dispersal is one type of movement that may be affected by TI^6–8^. This involves the movement of an individual from where it was born to a breeding patch (i.e. natal dispersal) or between successive breeding patches (i.e. breeding dispersal)^9,10^. Dispersal has major consequences on the demography of wild populations since dispersal rates affect the spatial synchrony of demographic parameters between patches and can reduce the risks of local extinction^11–12^. Dispersal also strongly affects the genetics of spatially structured populations by shaping genetic diversity and local adaptive processes^6,13^. There are two main effects TI can have on dispersers. First, mortality may occur due to collision with vehicles or stressful conditions during the crossing of the obstacle^2,4^: in this way, TI acts as a dispersal barrier. A second way TI can be a barrier to dispersal is by affecting an individual’s dispersal decisions (i.e. causing behavioral avoidance). Theoretical models have demonstrated that dispersal evolution is driven by the balance of costs and benefits of this behavior^9,10^. Roads and other obstacles may alter this cost–benefit balance by affecting, for instance, the energy expenditure or mortality risks related to crossing the obstacle^7^. Natural selection would be expected to favor behavioral avoidance of TI when an increase in dispersal costs resulting from TI outweighs dispersal benefits^7^.

Yet not all types of TI are permanent, impassable obstacles. The effects on mortality and dispersal decisions are expected to vary according to both the characteristics of the infrastructure and the intensity of the traffic^14^. In particular, one might expect mortality risk and behavioral avoidance to increase in line with the width of the TI, the artificiality of its surface, and traffic intensity^15,16^. Furthermore, individual factors could also influence mortality and dispersal behavior when organisms faced with TI (i.e. phenotype-dependent dispersal^9^). An individual’s tendency and capacity to disperse usually depend on a set of morphological, physiological, behavioral and life-history traits (i.e. dispersal syndrome^8,9,17^). Moreover, age also has a strong influence on dispersal propensity and ability^10,17^, which could result in TI having divergent effects on natal and breeding dispersal. One might expect, for example, that a juvenile’s lower body mass, inexperience and limited locomotive capacity would result in higher mortality and an enhanced dispersal cost in comparison to adults, which would increase avoidance behavior of TI before sexual maturity.

Although the effects of TI on dispersal have been studied in various taxa such as insects, mammals and birds^18–21^, no studies have simultaneously examined the effects of TI on natal and breeding dispersal, precluding any general evaluation of potential age-specific responses to TI. Yet this type of investigation is critical to better understand how TI affects the dispersal process during an organism’s ontogenesis and how it influences the dynamics of spatially structured populations. Age-dependent mortality during dispersal resulting from TI could have various consequences on long-term population viability, given that the sensitivity of the population growth rate to variation in juvenile and adult survival may differ between populations and species^22^. Moreover, as the relative contribution of natal and breeding dispersal rates and distances to dispersal dynamics as a whole may differ between populations and taxa^23^, TI could have a variety of effects on the viability of spatially structured populations.

In this study, we examined the effects of TI on dispersal at different ontogenetic stages in a pond-breeding amphibian, the yellow-bellied toad (*Bombina variegata*). This endangered species has recently become extinct in several regions of Western Europe^24^ and is therefore protected by national legislation over much of its range (it is registered in Appendix II of the Bern Convention and in Appendices II and IV of the EU Habitat Directive). We used capture-recapture data collected during a five-year period in a large spatially structured population of *B. variegata* (8,463 individuals captured in 189 breeding patches). We extended the multievent capture–recapture (CR) model developed by Lagrange et al.^25^ to examine the effects of two kinds of TI, gravel tracks and paved roads, on the dispersal of this species. First, we examined whether departure rates and dispersal distances differed between age classes. Next, we examined how age might affect an individual’s dispersal when faced with TI and tested the following predictions:

1. Molecular studies have reported that gravel tracks and paved roads may limit gene flow in amphibians^26,27^. Therefore, we expected a negative effect of TI on dispersal in the three age classes.
2. Paved roads are wider (resulting in greater clearance of vegetation), have an artificial surface, and support more intense traffic than gravel tracks, so might be expected to result in higher mortality and/or road avoidance behavior in dispersing organisms. We thus assumed that paved roads would have a higher negative effect on dispersal in all three age classes.
3. As body size positively influences locomotive capacity and reduces dispersal mortality in amphibians^28^, we predicted that the negative effect of TI on dispersal should decrease with age.

## Results

During the five-year study period, we captured 16,477 toads. Of these, we identified 8474 individuals: 2710 juveniles, 2158 subadults and 3606 adults (1805 males and 1801 females).

### Age-dependent departure rates and dispersal distances

We detected 265 dispersal events in juveniles, 268 dispersal events in subadults, and 507 dispersal events in adults. The raw distribution of dispersal distances for the three age classes is provided in Fig. 1. The median dispersal event distance was 695 m (coefficient of variation: 0.71) in juveniles, 568 m (CV: 0.792) in subadults, and 444 m (CV: 0.85) in adults. The distance covered during the longest dispersal events was 4529 m in juveniles, 3621 m in subadults, and 3141 m in adults.

**Fig 1.**
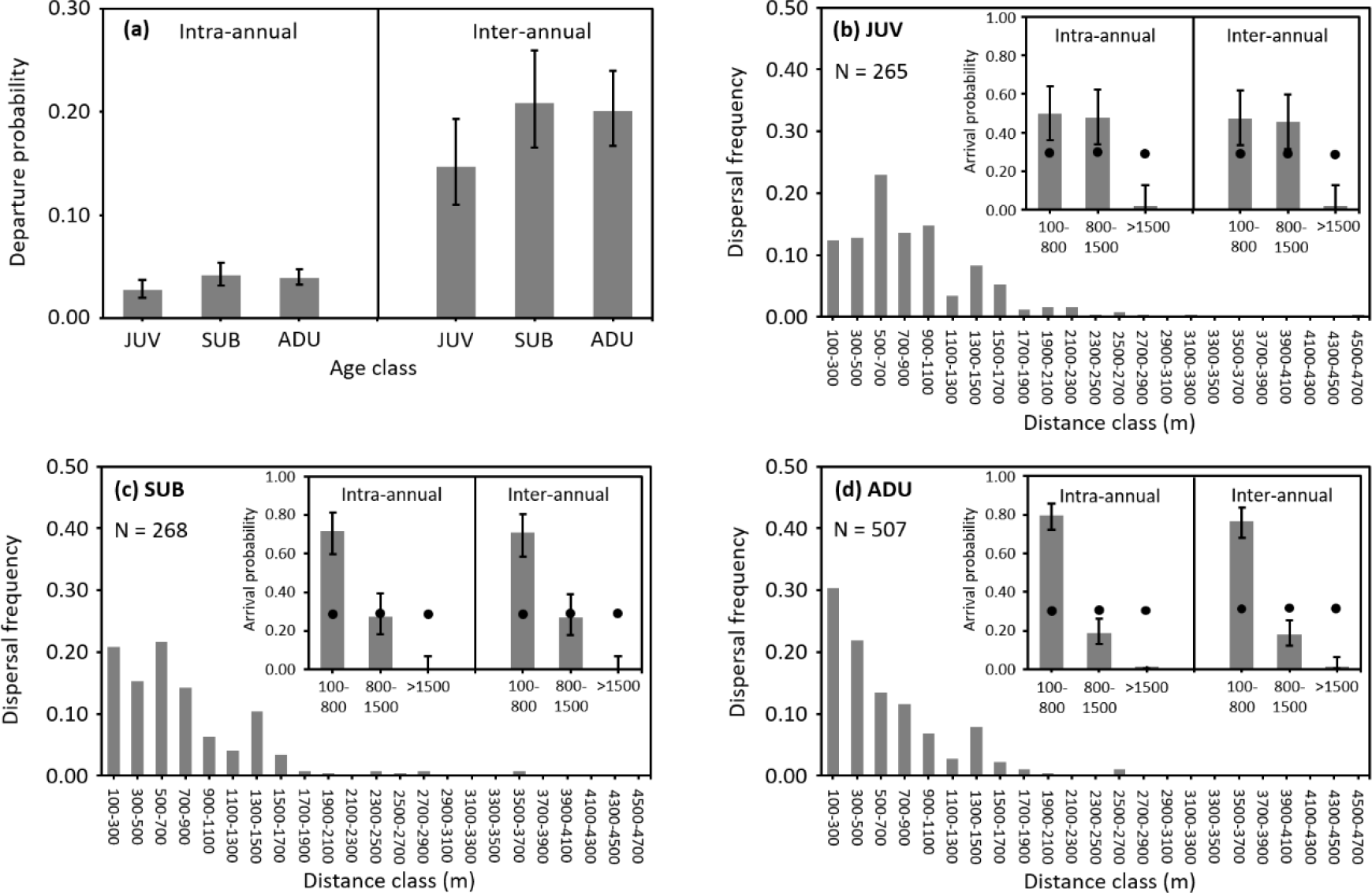
Age-dependent departure (a) and dispersal distance (b–d) in Bombina variegata. Three age classes are considered: juvenile (JUV, defined by one overwintering), subadult (SUB, two overwinterings) and adult (ADU, three or more overwinterings). Departure probability was estimated intra-annually (between two capture sessions within a year) and inter-annually (between two capture sessions in different years) in the multievent CR models. Concerning arrival and dispersal distances, the graphs show the (unmodeled) frequency of dispersal distances recorded for juveniles (b), subadults (c) and adults (d). Note that the movements do not necessarily occur between two consecutive capture sessions and can be intra-annual or inter-annual. The graphs also show the modeled probability (extracted from the multievent CR models) for an individual to arrive in a patch located at a distance of 100–800 m, 800–1500 m, and >1500 m from the source patch. The probability of an individual (whatever its age) arriving in a patch within each distance class is compared to the expected value, i.e. the probability calculated using the random dispersal hypothesis (shown as dots).

The best-supported model was [ϕ(AGE), ψ(AGE), α (AGE), *p*(AGE + YEAR)] (see model selection procedure in Supplementary Table S3). It indicated that recapture varied between age classes and years. In juveniles, recapture probability ranged from 0.14 (95% CI 0.07–0.27) in 2016 to 0.41 (95% CI 0.36–0.46) in 2012. In subadults, it ranged from 0.19 (95% CI 0.18–0.22) in 2015 to 0.31 (95% CI 0.28–0.35) in 2016. In adults, it ranged from 0.20 (95% CI 0.19–0.21) in 2015 to 0.37 (95% CI 0.35–0.38) in 2016. Survival also varied between age classes and increased with age. It was 0.46 (95% CI 0.43–0.49) in juveniles, 0.62 (95% CI 0.58–0.65) in subadults, and 0.66 (95% CI 0.64–0.68) in adults. Departure probability differed between age classes and was lower in juveniles than in subadults and adults (Fig.1). Our model also revealed that dispersal distance decreased with age (Fig.1). For the distance class 100–800 m, the deviation between the estimated and expected value was 25.00% higher in juveniles, 121.87% higher in subadults, and 146.87% higher in adults. For the distance class 800–1500 m, the deviation between the estimated and expected value was 70.59% higher in juveniles, 20.59% lower in subadults, and 44.11% lower in adults. For the distance class > 1500 m, the deviation between the estimated and expected value was 93.75% lower in juveniles, and 96.87% lower in subadults and adults.

### Age-dependent response to gravel tracks

In juvenile toads, we recorded 163 dispersal events without crossing a track and 115 dispersal events that involved crossing a track. In subadults, we recorded 172 dispersal events without crossing a track and 110 events that involved crossing a track. In adults, we recorded 324 dispersal events without crossing a track and 222 events that involved crossing a track. So movements that involved crossing a track represented 41% of juvenile dispersal events, 39% of subadult dispersal events, and 41% of adult dispersal events.

The best-supported model [ϕ(AGE), ψ(AGE), α (AGE), *p*(AGE + YEAR)] (see model selection procedure in Supplementary Table S4) showed that both survival and departure rates depended on age, and that recapture varied according to age and between years. The structure of this and the previous model is the same as regards survival and departure probabilities, so these estimates are not included again here. The model revealed that arrival probability was negatively affected by gravel tracks (Fig.2). At the intra-annual level, the proportion of individuals crossing a gravel tracks while dispersing was 0.22 (95% CI 0.12–0.36) in juveniles, 0.29 (95% CI 0.18–0.42) in subadults, and 0.31 (95% CI 0.22–0.40) in adults. At the inter-annual level, we detected a similar pattern: the proportion of individuals crossing an unpaved track while dispersing was 0.36 (95% CI 0.22–0.52) in juveniles, 0.44 (95% CI 0.32–0.57) in subadults, and 0.46 (95% CI 0.35–0.58) in adults. These values are systematically far lower than those expected based on a random dispersal hypothesis: 0.89 in juveniles and 0.90 in adults and subadults. The deviation of the estimated from the expected value was 75.28% lower in juveniles, 67.77% lower in subadults, and 65.55% lower in adults at the intra-annual level; at the inter-annual level, the deviation was 59.55% lower in juveniles, 51.11% lower in subadults, and 48.89% lower in adults. Overall, our analyses revealed that gravel tracks had a negative effect on dispersal and that juveniles were slightly more sensitive to these tracks than subadults or adults.

**Fig 2.**
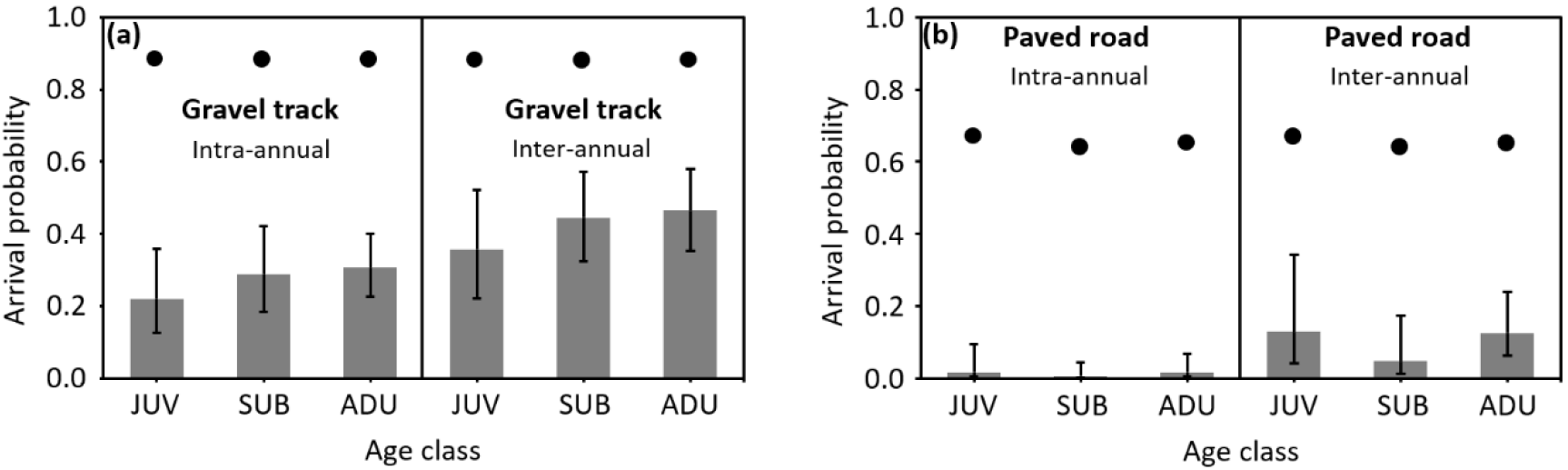
Influence of gravel tracks and paved roads on age-dependent dispersal in Bombina variegata. Three age classes are considered: juvenile (JUV, defined by one overwintering), subadult (SUB, two overwinterings) and adult (ADU, three or more overwinterings). The probability of an individual (whatever its age) arriving in a patch after crossing a gravel track (a) or a paved road (b) is systematically lower than the expected value, i.e. the probability calculated using the random dispersal hypothesis (shown as dots). The 95% confidence intervals are shown as error bars.

### Age-dependent response to paved roads

In juveniles, we recorded 255 dispersal events without crossing a road and only 23 dispersal events that involved crossing a road. In subadults, we detected 269 dispersal events without crossing a road and 13 events that involved crossing a road. In adults, we recorded 511 dispersal events without crossing a road and 35 events that involved crossing a road. So, movements that involved crossing a paved road represented 9% of juvenile dispersal events, 5% of subadult dispersal events, and 6% of adult dispersal events.

The best-supported model [ϕ(AGE), ψ(AGE), α (AGE), *p*(AGE + YEAR)] (see model selection procedure in Supplementary Table S5) indicated that arrival probability was negatively influenced by paved roads (Fig.2). At the intra-annual level, the proportion of dispersers that crossed a road was 0.01 (95% CI 0.003–0.09), 0.01 (95% CI 0.0007–0.04) in subadults, and 0.02 (95% CI 0.004–0.07) in juveniles and adults. At the inter-annual level, the proportion of dispersers that crossed a road was 0.13 (95% CI 0.04–0.34) in juveniles, 0.05 (95% CI 0.01–0.17) in subadults, and 0.12 in adults (95% CI 0.06–0.24). These values are drastically lower than those expected based on a random dispersal hypothesis, which predicts the probability of crossing a paved road as 0.67 in juveniles, 0.64 in subadults and 0.65 in adults. The deviation of the estimated from the expected value was 97.46% lower in juveniles, 99.06% lower in subadults, and 97.38% lower in adults at the intra-annual level; at the inter-annual level, the deviation was 80.59% lower in juveniles, 92.19% lower in subadults, and 81.53% lower in adults. Moreover, our results found that paved roads have a higher negative effect on dispersal than gravel tracks. At the intra-annual level, the deviation of the estimated from the expected value was always higher (1.29 times in juveniles, 1.46 times in subadults, and 1.48 times in adults) for roads than for tracks. Similarly, at the inter-annual level, the deviation of the estimated from the expected value was also systematically higher (1.35 times in juveniles, 1.80 times in subadults, and 1.67 times in adults) for roads than for tracks. Overall, our analyses showed that paved roads have a strong negative effect on dispersal and that all three age classes are impacted in a relatively similar way.

## Discussion

Our study found generally lower emigration rates for juveniles than for subadults and adults. It also revealed that dispersal distance decreased with age. Additionally, it highlighted that both tracks and roads have negative effects on dispersal, but the negative effect of paved roads was higher than that of gravel tracks. The effect of tracks on dispersal slightly decreased with age; in contrast, the negative effect of paved roads was severe and relatively similar across the three age classes.

Our results showed emigration rates were high overall (between 0.03 and 0.04 intra-annually and between 0.15 and 0.21 inter-annually) in the three age classes compared to that reported for other amphibian species^29^. This low site fidelity is due to the low inter-annual persistence of this species’ breeding patches in forest environments^30,31^. A recent study has demonstrated that high patch turnover in harvested woodlands led to high dispersal rates in *B. variegata*^30^. Our findings also highlighted age-dependent dispersal rates and distances. At the juvenile stage, individuals were less likely to emigrate than at subadult and adult stages. Yet when they decided to emigrate, juveniles and subadults undertook longer dispersal distances than adults. These results are congruent with the idea that emigration decisions depend on phenotypic characteristics^32^. In amphibians, a juvenile’s body size is thought to influence the decision-making process that initiates emigration from the natal pond^28^. It is likely that juveniles need to attain a certain size threshold to make the decision to disperse, which results in lower emigration rates at this stage. However, after leaving their natal patch, the juveniles and subadults in our study performed larger dispersive movements than adults. The largest dispersal events were undertaken by a juvenile (4529 m) and a subadult (3621 m), both greater than the largest distance covered by an adult (3141 m). Interestingly, the distribution of subadult dispersal distances was intermediate – between that of juveniles and adults – which suggests a progressive behavioral shift during the period preceding first reproduction. This progressive change in dispersal behavior might be the result of a shift in the causal factors driving dispersal distances. In particular, dispersing farther before sexual maturity could allow individuals to mitigate the risk of kin competition and inbreeding^10,32,33^. After sexual maturity, breeding dispersal is more likely to be influenced by environmental conditions (e.g. the quantity and quality of resources) in breeding patches^10^. In this context, dispersing long distances is not really necessary when suitable patches occur in the vicinity of the patch currently occupied.

Our results validated our prediction about the effects of TI: tracks and roads have a negative impact on dispersal at all ages. This detrimental effect can be caused by direct mortality or by behavioral avoidance. Our analyses revealed relatively low survival rates in the three age classes compared to those reported in other populations of the studied species^31^. This decrease in survival could potentially result from over-mortality induced by TI. Collision with vehicles may increase mortality rates; indeed, studies have reported massive mortality caused by collision during migration in a broad range of amphibian taxa^34^. However, we did not find evidence of massive mortality on either tracks or roads in our study – although mortality resulting from crossing TI during dispersal would not be easily detectable (due to the small number of carcasses spread over large distances, rapid disappearance of carcasses, etc.). Moreover, on the gravel tracks, vehicle traffic is very low (less than one vehicle per day), and is also relatively low on the paved roads (between 70 and 80 vehicles per day), making this an unlikely source of mortality. If an over-mortality occurs, it would be rather caused by the loss of vegetation due to the presence of TI. For amphibians, the absence of vegetation increases the level of evapotranspiration and thus reduces their chances of survival during dispersal^35,36^. Additionally, the negative effect of TI on natal dispersal and breeding dispersal found in our study could result from behavioral avoidance of tracks and roads. Studies have reported that amphibians are sensitive to ground surfaces when they move^37,38^ and avoid surfaces without vegetation cover^35^. In this way, TI could act as a dispersal barrier through two non-mutually exclusive mechanisms: over-mortality caused by dehydration during TI crossings and/or behavioral avoidance when faced with the mortality risks associated with these crossings.

Our findings validated our prediction about the impact of differences in TI permeability (that is, permeable for amphibian crossings): paved roads had a more detrimental impact on dispersal than gravel tracks. This pattern could result from differences in TI – in particular, it is possible that the artificiality of the TI surface affects *B. variegata* behavioral response. In our study area, tracks are surfaced with compressed conglomerates, which are relatively natural substrates. In contrast, roads paved with asphalt and coal tar contain complex mixtures of volatile and non-volatile chemical compounds that could elicit road-avoidance behavior. Amphibians have a highly developed olfactory sense, which plays a critical role in their movements and orientation^39,40^. Studies have reported that anurans are highly sensitive to the composition of the ground cover when they move^37,38,41^ and tend to avoid asphalted areas^42^. Additionally, the width of TI could also explain the divergent effects of tracks and roads on dispersal. Wider TI automatically result in a larger unvegetated surface area. Studies have reported that amphibians usually avoid moving across unvegetated or little vegetated ground in order to avoid mortality risks due to dehydration^35,37,38^.

Contrary to our expectations, our findings revealed that age has little effect on the response of *B. variegata* to TI. The negative effect of tracks appears to slightly decrease with age, with juveniles more impacted than subadults or adults. These results suggest that gravel tracks are average-permeability barriers whose negative effects may slightly decrease during ontogenesis. This may result from a decrease in mortality during the crossing of tracks and/or from a higher propensity to cross this kind of barrier with an increase in age or size. Studies have shown that amphibian mortality rates during dispersal (caused by desiccation, exhaustion and predation) decrease with body size^28^, and therefore age. Other works have also demonstrated that amphibians’ propensity for exploration and locomotive capacity increase with body size in juveniles^43,44^. In terms of paved roads, our results showed that dispersal across all three age classes was strongly and equally impacted by this type of TI. Paved roads are low-permeability barriers that seriously impact dispersal regardless of age. An increase in body size, experience and locomotive capacity would probably not allow an individual to mitigate the dispersal cost induced by roads over its lifetime.

To our knowledge, this study is the first attempt to evaluate the effects of TI on both natal and breeding dispersal at the landscape level in a vertebrate. It also emphasizes that TI with relatively small width and low traffic may act as important dispersal barriers in small-sized vertebrates both before and after sexual maturity. Moreover, the results highlight the importance of examining how factors related to the individual characteristics of an organism as well as the specific characteristics of TI may affect dispersal patterns. They suggest that age may affect dispersal decisions when individuals are faced to TI with small or moderate permeability. Yet, our study also demonstrates that enhanced dispersal costs induced by several TI cannot be mitigated during organism’s ontogenesis. From a conservation point of view, the determination of conservation measures should be based on a deeper understanding of the effects of TI on cost and benefits balance of dispersal. However, our knowledge about such effects remains to date very patchy in a broad range of taxa. In the context of a global intensification of transport networks, future studies should be undertaken to quantify more accurately the effects of different types of TI on individual dispersal strategies and the long-term dynamics of spatially structured populations.

## Materials and methods

### Study area and capture–recapture data

The study was conducted on a spatially structured population of *B. variegata* between 2012 and 2016 in northeast France (N 49° 11′ 44″, E 5° 26′ 00″) in the Verdun National Forest, which covers a surface area of 9,600 ha and consists of deciduous and coniferous trees. This population is divided into subpopulations that occupy aggregations of ruts and residual puddles resulting from logging activities – these are the toads’ breeding areas. Based on the method used in a previous study^30^, the breeding patches were delineated using QGIS by creating polygons connecting the waterbodies located on the boundary of the archipelago (the ‘Minimum Convex Polygon’ approach^44^). Breeding patches were assumed to be different when the minimal distance separating the boundary polygons exceeded 100 m. This distance was chosen since the movement distance between ponds in *B. variegata* is usually less than 100 m^46,47^. This allowed us to assume that the movements recorded between patches were more likely to correspond to dispersal events rather than routine, short-distance, inter-pond movements. This approach resulted in 189 distinct patches (Fig.3). The mean number of ponds per breeding patch was 4±3 (min. = 1, max. = 15). The median Euclidian distance between patches was 4,548 m (min. = 100 m; max. = 12,357 m).

**Fig 3.**
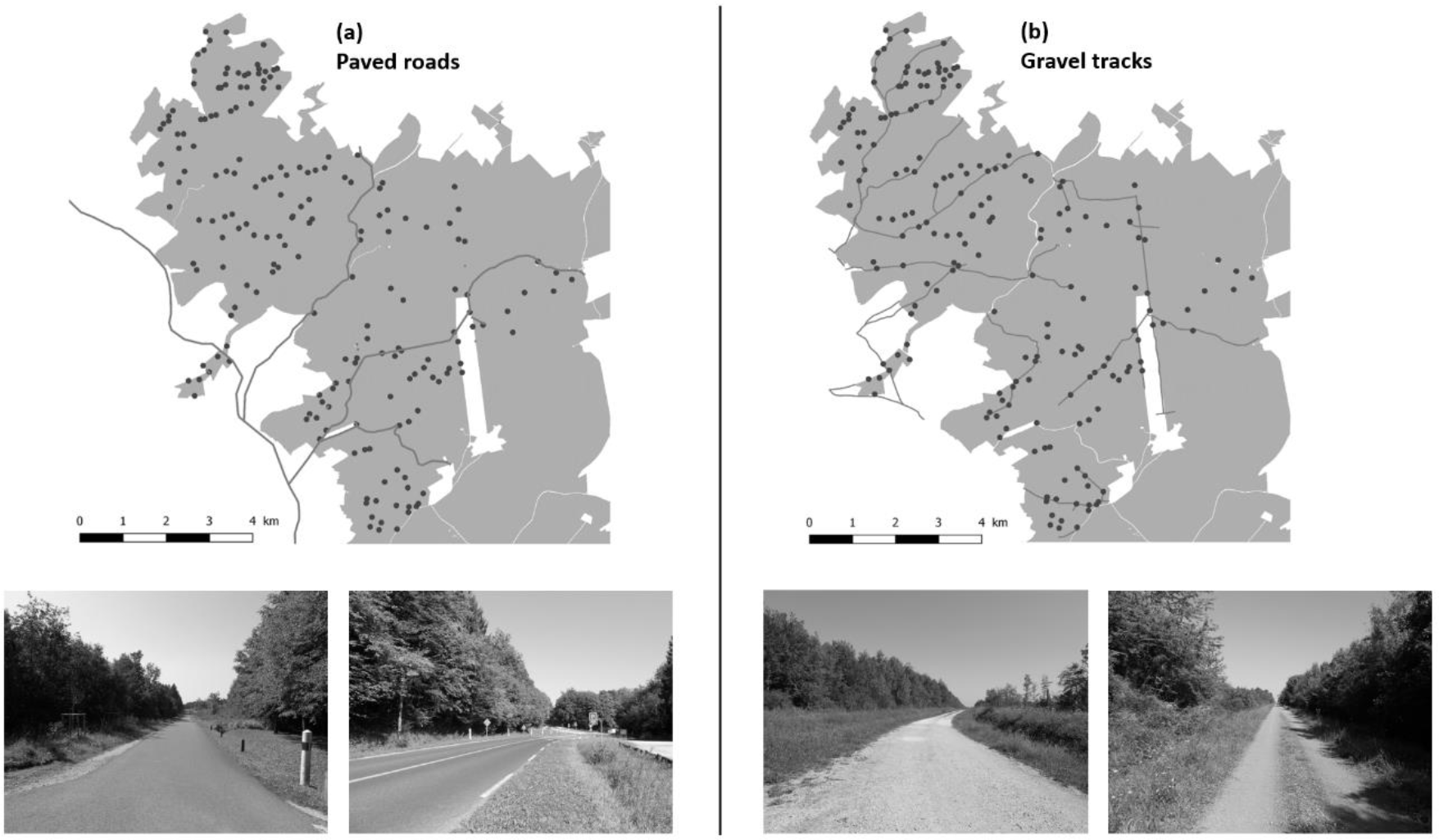
Map of study area showing the forest cover (grey background), the 189 B variegata breeding patches (black dots), the four paved roads (a: grey lines) and the 13 gravel tracks (b: grey lines).

The study area (Fig.3) encompassed 13 gravel tracks used by private vehicles (i.e. cars and motorbikes) and logging vehicles (i.e. heavy trucks and skidders) and 4 paved roads used by a large variety of vehicles (cars, heavy trucks, farming tractors, etc.). The two types of TI differ in terms of width, surface material and traffic intensity. Paved roads are wider (7–12 m) than gravel tracks (3.5–6 m), implying more vegetation clearance. The artificiality of the surface is also higher for paved roads (asphalt) than for gravel tracks (compressed aggregates). The traffic intensity is also higher on paved roads (between 70 and 80 vehicles per day; information from the Road Service of the Meuse department) than on gravel tracks (less than one vehicle per day).

Three capture–recapture sessions (in May, June and July) were carried out each year. During each capture session, we sampled each waterbody in the breeding patches to determine the presence of *B. variegata*. The toads were captured by hand or with a dipnet, and were then photographed and released back into the pond where they were caught. As in previous studies^30,48^, we considered three age classes: juveniles captured after their first overwintering, subadults (two overwinterings), and sexually mature adults (at least three overwinterings). Toad size (snout–vent length) ranged from 17 to 29 mm in juveniles and from 30 to 35 mm in subadults. We assumed that individuals become sexually mature at the age of 3^48^, with a mean body length of 36 mm in males and 37 mm in females. We identified each individual by the specific pattern of black and yellow mottles on its belly, recorded by photographs. Multiple comparisons of patterns were performed using a robust computing tool (Extract Compare) to minimize misidentification errors^49^.

Toad capture was authorized by the Préfecture de la Meuse (arrêté no. 2008–2150). We confirmed that toad capture was performed in accordance with relevant guidelines and regulations.

### Modeling age-dependent departure rates and dispersal distances

We modeled dispersal using multievent CR models^50^. Lagrange et al.^25^ proposed a model that allows the estimation of survival (ϕ) and dispersal (ψ) between numerous sites. By omitting site identity and distinguishing ‘individuals that stay’ and ‘individuals that move’, this model circumvents the computational issues usually encountered in classical multisite CR models when the number of sites is large. Lagrange’s model is based on states that embed information on whether an individual at *t* is occupying the same site as it occupied at *t*−1 (‘S’ for ‘stay’) or not (‘M’ for ‘move’), with information about whether the individual was captured (‘+’) or not (‘o’) at *t*−1 and at *t*. In a recent paper, Tournier et al.^51^ extended Lagrange’s model by breaking down dispersal (ψ) into the distinct parameters of departure (ε) and arrival (α). This new parameterization allows the quantification of the proportion of individuals arriving in sites of different characteristics, located at different distances from the source site, or crossing different types of obstacles.

We adapted this parameterization for the needs of our study. We considered states incorporating information about the capture of an individual (‘+’ or ‘o’) at *t*−1 and *t* and its movement status. We also included states with information about the age class of an individual: juvenile (‘j’), subadult (‘s’), and adult (‘a’). In addition, we included information about the Euclidian distance covered by dispersers between a departure and an arrival patch; this was incorporated in the model using three distance classes: 100–800 m (‘1’), 800–1500 m (‘2’), > 1500 m (‘3’). This led to the consideration of 37 states in the model (Supplementary Table S1). For instance, an individual +jS+ was captured at *t*−1 and at *t*, is in the juvenile class, and remained in the same patch between *t*-1 and *t*. An individual +sM1+ was captured at *t*−1 and at *t*, is in the subadult class, does not occupy the same patch as at *t*−1, and arrived in a patch at a distance of between 100 m and 800 m from the source patch. We distinguished between a total of 16 events, which are coded in an individual’s capture history and reflect the information available to the observer at the time of capture (Supplementary, Table S2). The models had a robust design structure^52^, i.e. the three capture sessions performed each year corresponded to secondary sessions, and each set of three sessions corresponded to a primary period. This robust design structure allowed us to examine both intra-annual and inter-annual dispersal.

When captured for the first time, the state of an individual could be ojS+, osS+ or oaS+. We then considered five steps during which the information of the state descriptor was progressively updated in the model: survival (ϕ), departure (ε), arrival (α), age transition (δ) and recapture (*p*). Each step was conditional on all previous steps. As in Lagrange et al.^25^ and Cayuela et al.^53^, in the transition matrix (Fig.2 and Fig.3), the rows correspond to time *t*−1, the columns to time *t*, and whenever a status element is updated to its situation at *t*, it is shown in bold and stays bold throughout the following steps. In the first step, the information about survival was updated. An individual could survive with a probability of ϕ or could die (**D**) with a probability of 1–ϕ. This led to a matrix with 37 states of departure and 7 intermediate states of arrival (Fig.4). As the time period between two secondary sessions was shorter (1 month) compared to the lifetime of an individual of this species^30^, survival probability was fixed at 1 between secondary sessions. In the second modeling step, departure was updated. An individual could move (**M**) from the site it occupied with a probability of ε or could stay (**S**) with a probability of 1–ε. This led to a matrix of 7 states of departure and 13 states of arrival (Fig.4). In the third step, we updated the information about arrival. An individual that moved could arrive in a patch located in the first two distance classes (**1** or **2**) from the source patch with a probability of α, or could arrive in a site located in distance class (**3**) with a probability of 1–α. This led to a matrix with 13 states of departure and 25 states of arrival (Fig.4). In the fourth modeling step, information about the individual’s age was updated. Between secondary sessions, an individual was forced to stay in its age class (**j**, **s** or **a**) with a probability of δ = 1. Between primary sessions, an individual was constrained to reach the following age class with a probability of δ = 1. This led to a transition matrix with 25 states of departure and arrival (Fig.4). Note that the individuals belonging to the adult age class (**a**) were forced to stay in their class. In the fifth and last step, recapture was updated (Fig.5). An individual could be recaptured with a probability of *p* or not recaptured with a probability of 1–*p*, resulting in a transition matrix with 25 states of departure and 37 states of arrival. The last component of the model linked events to states. In this specific situation, each state corresponded to only one possible event (Fig.5).

**Fig 4.**
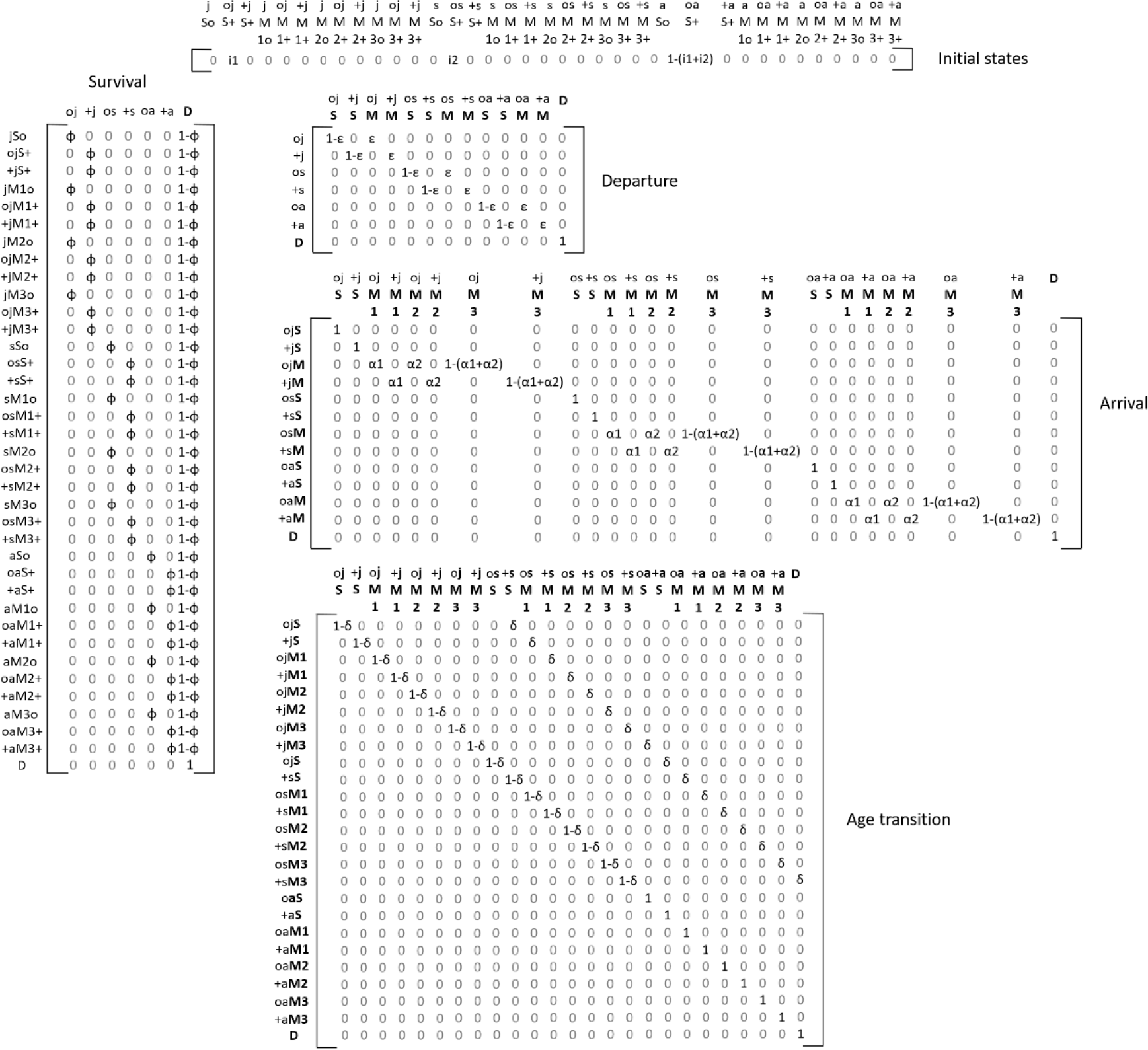
Matrices for the initial states of departure and the four first successive modeling steps: survival, departure (from patch), arrival (to patch) and age transition. The 37 states of the model are described in Supplementary, Table S1. The information updated at each step appears in bold.

**Fig 5.**
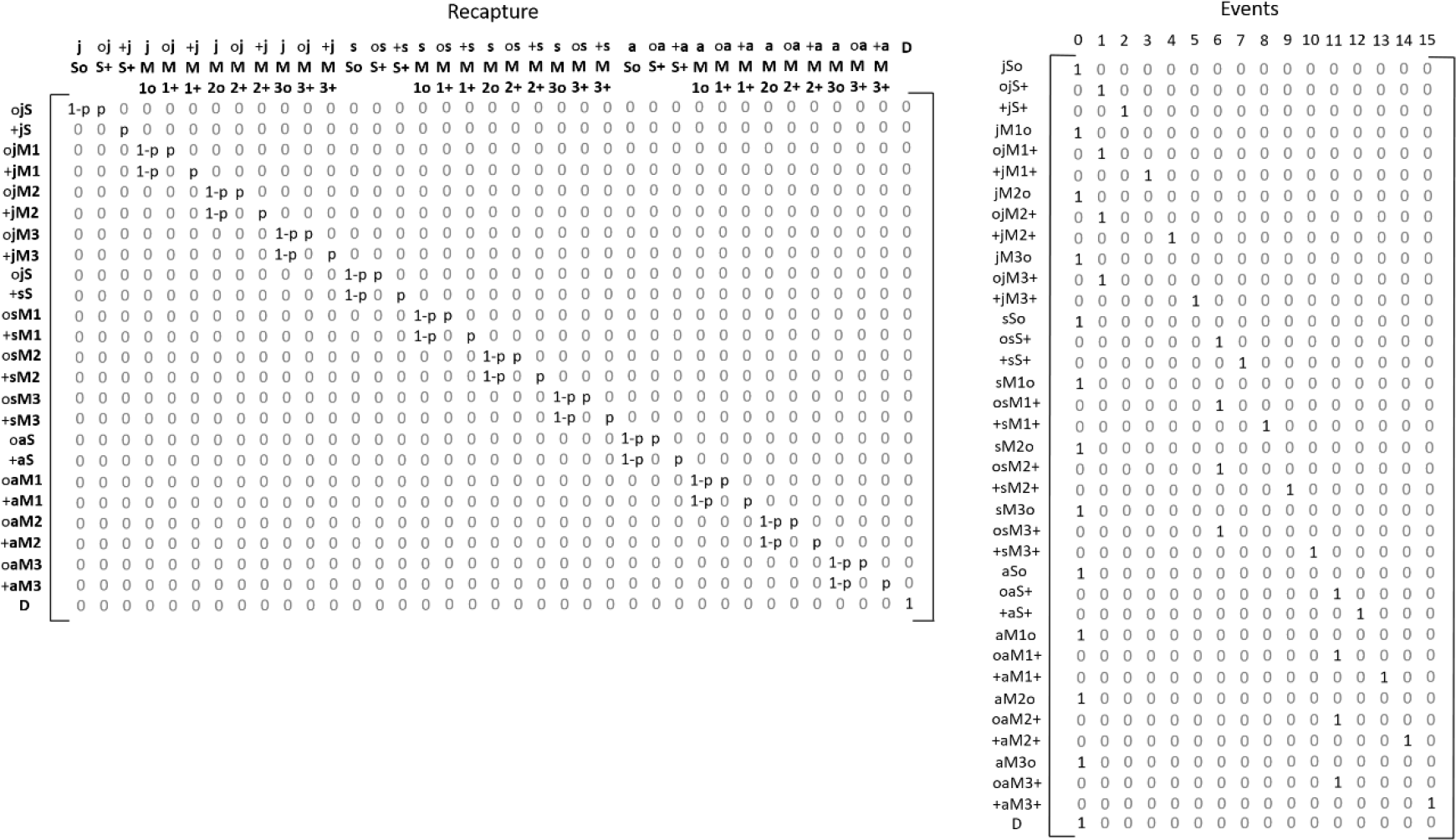
Matrices of the fifth modeling step (i.e. recapture) and of events (field observations). The 37 states and events of the model are described in Supplementary, Table S1 and Table S2 respectively.

This parameterization was implemented in the E-surge program^54^. Competing models were ranked through a model selection procedure using Akaike information criteria adjusted for a small sample size (AICc) and AICc weights. We considered that two models were differently supported by the data when the ΔAICc was > 2. Our hypotheses concerning recapture and state-state transition probabilities were tested from the general model [ϕ(AGE), ε(AGE), α(AGE), δ(.), *p*(AGE + Y)], which includes two effects: (1) the three age classes (AGE) coded as states in the model, and (2) year-specific variation (Y). The notation (.) means that a parameter is held constant. From this general model, we tested all the possible combinations of effects and ran 16 competitive models (Supplementary Table S3).

### Modeling age-dependent response to gravel tracks and paved roads

To model the effect of gravel tracks and paved roads on *B. variegata* dispersal, we used a restricted version of the model described above (28 states and 13 events). This model had the same architecture as the one previously described, but the three distance classes were replaced by two classes related to crossing TI: individuals could arrive in a new patch (1) without crossing any track or road, or (2) by crossing one (or more) track or road. To avoid difficulties in model implementation, the effects of tracks and roads were analyzed separately. The model selection was performed in a similar way as previously described for age-dependent dispersal distances. Sixteen competitive models were run to test the effects of both types of transport infrastructure (Supplementary Table S4 and Table S5).

### Assessing the effects of Euclidean distance and TI on dispersal

The conditional probability of arrival (i.e. depending on the Euclidean distance between breeding patches) estimated by the multievent CR models strongly depended on the quantity of patches within each distance class and the number of individuals in each patch that may reach these. Similarly, arrival probability also strongly depended on the quantity of TI in the area and the number of individuals that may cross these.

For this reason, we examined the effects of Euclidean distance and TI by comparing the conditional arrival probability extracted from the best-supported models to the probability of reaching a patch in each distance class using a random dispersal hypothesis (i.e. the mean probability of arriving in a patch calculated from all the individuals occurring in all patches of the study area). In addition, to assess the effects of TI, we compared the arrival probability provided by the best-supported model to the probability of crossing a track or a road using a random dispersal hypothesis (i.e. the mean probability of crossing a track or road calculated from all the individuals occurring in all patches of the study area). As the number of juveniles, subadults and adults varied among patches, we calculated the expected random age-specific probability. We assumed that the effects of Euclidean distance and TI were significant if the 95% CI of the conditional arrival probability did not overlap the expected random probability. The percentage of deviation from the expected random probability was used to assess the influence of Euclidean distance and to rank the effect of tracks and roads on dispersal and between age classes.

## Acknowledgments

We would like to warmly thank all the fieldworkers who helped in data collection. We are also grateful to the French National Office of Forests for its financial and technical support.

## Author Contributions

HC, AB and EB conceived and designed the study. EB performed the study. HC analyzed the data. HC and AB wrote the manuscript.

## Additional information

Supplementary information accompanies this paper at …

